# GAPDH is tethered to axonal transport vesicles by S-acylation

**DOI:** 10.64898/2026.09.17.752404

**Authors:** Nisandi N. Herath, Jordan A. Kogut, Andrey A. Petropavlovskiy, Amelia H. Doerksen, Charlotte A. Townsend Bennie, Anthony Dang, Gareth M. Thomas, Dale D.O. Martin, Shaun S. Sanders

**Author notes:** These authors contributed equally (Nisandi N. Herath, Jordan A. Kogut).

## Abstract

Vesicle movement along axonal microtubules in neurons requires the ATP-dependent molecular motors dynein and kinesin. Fast axonal transport is fueled by ATP, provided by vesicle-associated glycolytic enzymes, but how these predicted soluble enzymes attach to vesicles in unclear. One potential mechanism is the protein lipid modification S-acylation, which involves the addition of long chain fatty acids to protein cysteine residues mediated by the ZDHHC (Asp–His–His–Cys) family of protein S-acyltransferases. Among the many effects this lipid modification imparts is an increase in protein localization to membranes. We found that eight of the ten glycolytic enzymes are S-acylated in the brain. Of the 10 glycolytic enzymes, we focused on glyceraldehyde 3-phosphate dehydrogenase (GAPDH) as it is the first enzyme of the payoff phase of glycolysis. GAPDH is S-acylated on cysteine 247 by ZDHHC5 and ZDHHC17. Importantly, C247 point mutation impairs GAPDH association with vesicles in hippocampal neurons. Investigating the role of S-acylation in glycolytic enzyme localization will lead to novel insights into neuronal transport mechanisms and may also shed light on neurodegenerative disease pathology and potential drug targets.

## Introduction

Neurons are large, polarized cells with output projections called axons that send signals to downstream targets. Axons can be very long, up to a meter in corticospinal tract and the sciatic nerve, and, as such, require efficient delivery of cargo to and from distal sites^1^. Axonal transport is critical for neuronal function with impaired transport implicated in various neurodegenerative and neurological disorders, including Huntington, Alzheimer, and Parkinson diseases, amyotrophic lateral sclerosis, spinal muscular atrophy and various paraplegias and neuropathies^2–5^. Thus, understanding the mechanisms that govern axonal transport may provide insight into how disrupted axonal transport contributes to neuronal dysfunction and disease progression.

Membrane-bound cargo such as endosomes, lysosomes, and synaptic vesicles are trafficked by the molecular motor ATPases (adenosine triphosphatases) dynein and kinesin along axonal microtubules at speeds reaching 400 mm/day^6–9^. Kinesin motors transport cargo in the anterograde direction, toward microtubule plus ends at axon terminals, while dynein traffics cargo in the retrograde direction towards microtubule minus ends at the soma^8–12^. Both motor proteins require ATP to fuel each 8 nm step; therefore, a constant, local energy source is critical to maintain fast axonal transport^13–15^. ATP produced by a population of vesicle-bound glycolytic enzymes, is necessary and sufficient to meet the energy demands of molecular motors during fast axonal transport of non-mitochondrial cargo^16,17^.

How glycolytic enzymes tether to fast-moving vesicles to facilitate transport is not fully understood. Wild type (WT) Huntingtin (HTT), the protein mutated in Huntington disease (HD), plays a role in scaffolding glyceraldehyde 3-phosphate dehydrogenase (GAPDH) on vesicles.

However, the impact of loss of HTT on fast axonal transport was modest, so clearly other factors play a role^17^. One potential mechanism is the protein lipid modification S-acylation (commonly palmitoylation). S-acylation is the reversible addition of long-chain fatty acids to cysteine residues via a labile thioester bond to regulate protein localization and function. For soluble cytosolic proteins, like the glycolytic enzymes, the hydrophobic fatty acyl chain can provide a tether to phospholipid bilayers^18^. Interestingly, proteomic studies suggest that all glycolytic enzymes are S-acylated in several S-acyl-proteomic studies, and GAPDH can be S-acylated when treated with palmitoyl-CoA *in vitro*^19–23^. However, GAPDH S-acylation in intact cells has not been shown, and the functional consequence of this modification is also unknown.

Thus, we sought to confirm S-acylation of the glycolytic enzymes in the nervous system using low throughput methods. Focusing on the key initial enzyme of the payoff phase of glycolysis, GAPDH, we also sought to determine if S-acylation serves as a mechanism to tether GAPDH to vesicles in neurons.

## Results

### Six glycolytic enzymes are S-acylated predominantly in the nervous system

All glycolytic enzymes have been identified in at least 15 S-acyl proteomic studies from human, mouse and rat^19^. As only HK, GAPDH, PK have been verified as S-acylated in low-throughput experiments^19–23^, we first sought to determine if all the glycolytic enzymes are indeed S-acylated *in vivo*. To do so we purified S-acylated proteins from brain, heart, and kidney of female Sprague-Dawley rats using the acyl biotin exchange (ABE) assay. Indeed, eight out of ten glycolytic enzymes are S-acylated (HK, glucose 6-phosphate isomerase [GPI], phosphofructokinase [PFK], aldolase [ALDO], triosephosphate isomerase [TPI], GAPDH, phosphoglycerate kinase [PGK], and PK) and six are S-acylated significantly more in the brain compared to the heart and kidney (HK, GPI, PFK, GAPDH, PGK, and PK; Figure 1). S-acylation of phosphoglycerate mutase (PGAM) or enolase (ENOL) was not detected in any tissue examined (Figure 1). In contrast, S-acylation of calnexin (CALX), a known S-acylated protein used as a positive control, was readily detected in all three tissues (Figure 1). Importantly, CREB (cAMP-response element binding protein), a non-S-acylated protein used as a negative control, was not detected in ABE samples, confirming the specificity of the assay (Figure 1).

**Figure 1.**
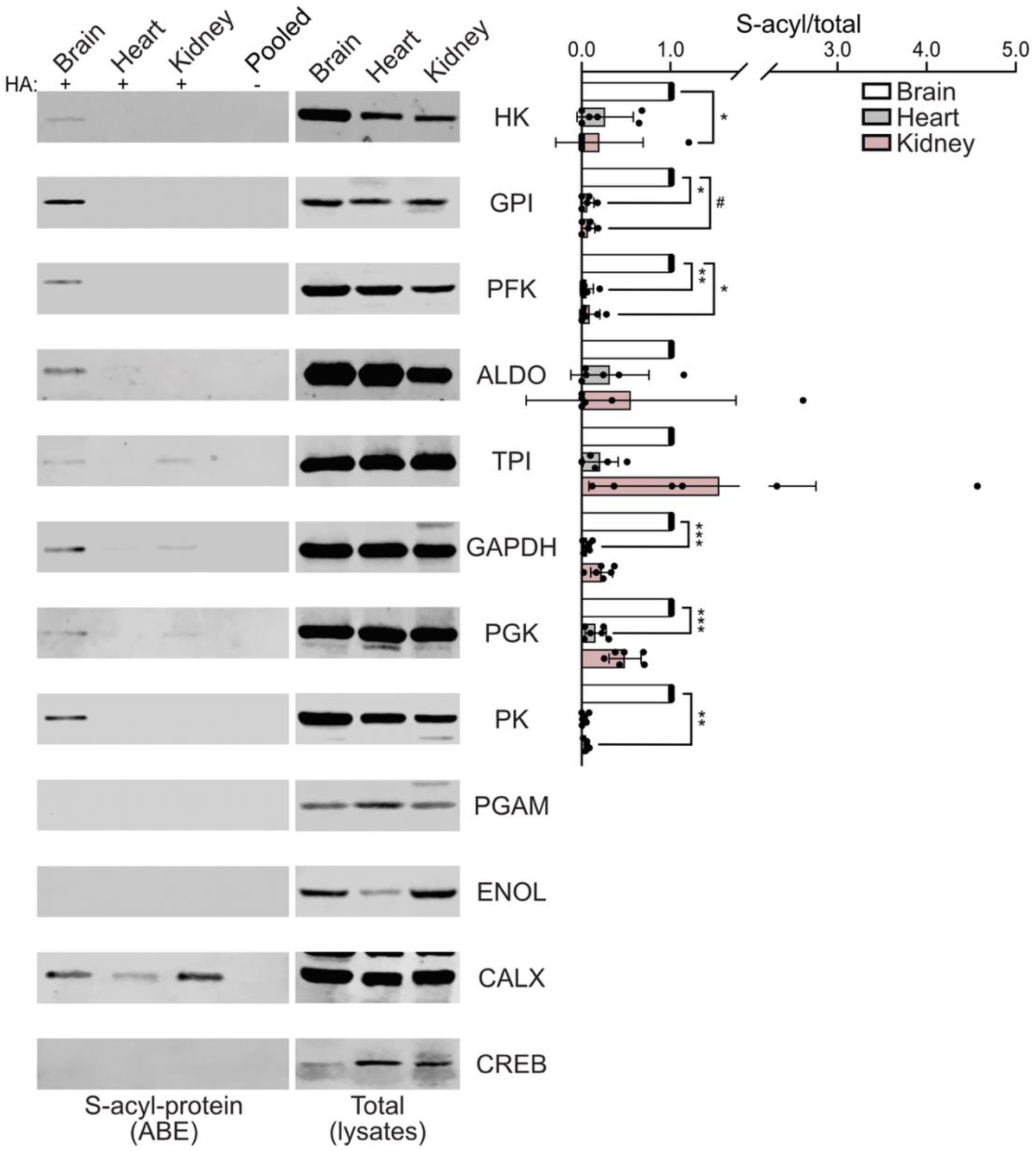
Eight of ten glycolytic enzymes are S-acylated *in vivo* and six are more S-acylated in nervous tissue. Rat tissues were lysed and S-acyl-protein levels (isolated by ABE; left panels) and total protein levels in parent lysates (right panels) were assessed by immunoblot with antibodies against the indicated glycolytic enzymes and control proteins (CALX, positive control, and CREB, negative control). Quantified data are shown on the right with S-acyl/total levels, normalized to the brain (Kruskal-Wallis test, N=6; HK: p=0.0058, Kruskal-Wallis statistic [KW]=8.92, *p=0.016; GPI: p=0.0018, KW=9.925, *p=0.012, #p=0.034; PFK: p=0.00040, KW=11.84, **p=0.0061, *p=0.013; GAPDK: p<0.0001, KW=14.17, ***p=0.0005; PGK: p<0.0001, KW=15.30, ***p=0.003; PK: p<0.0001, KW=12.45, **p=0.0017). Pooled is a combined sample processed without the key ABE reagent, hydroxylamine (HA; negative control).

### PK, PGK, and GAPDH are S-acylated in HAP1 cells

Our findings in Figure 1 demonstrate that a subset of the glycolytic enzymes are S-acylated *in vivo*. However, while proteins can be modified with various long-chain fatty acids^24^, the ABE assay does not discriminate the fatty acid or fatty acids each protein is modified with. Instead, assessing S-acylation in cultured cells using bioorthogonal labeling with various fatty acids followed by purification by click chemistry can provide insight into the fatty acid preferences for a given protein. In the brain, 16- and 18-carbon fatty acids, saturated and unsaturated, are the most commonly used for S-acylation^24^. We therefore assessed GAPDH, PK, and PGK S-acylation using bioorthogonal labeling and click chemistry with alkyne palmitate and alkyne stearate in HAP1 (human haploid) cells, a cell type in which glycolytic enzyme S-acylation was previously detected by S-acyl proteomics^25^. All three enzymes are readily modified with both fatty acids, although there are non-significant trends towards preferences for palmitate over stearate for PGK and PK (Figure 2).

**Figure 2.**
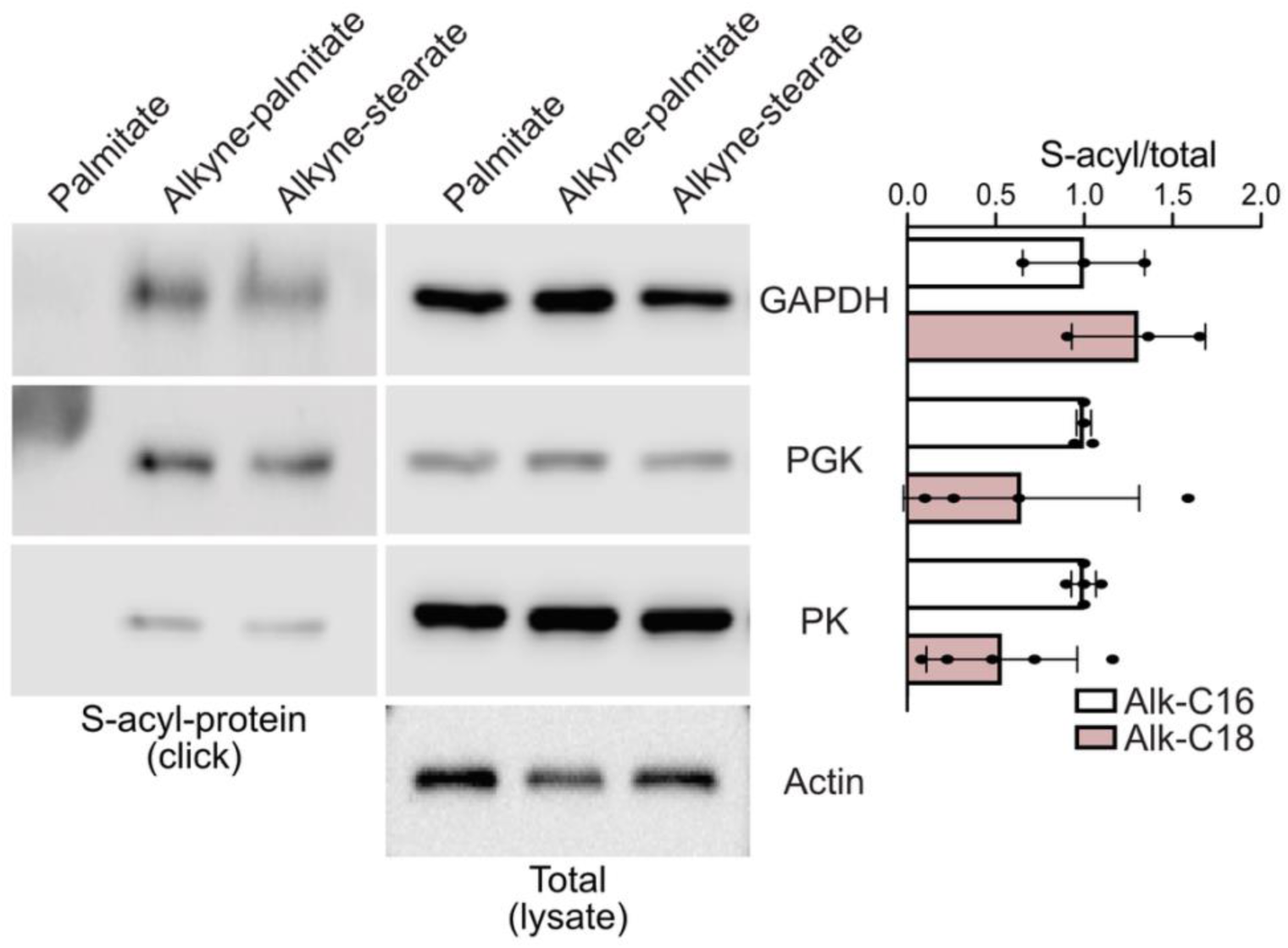
GAPDH, PGK, and PK are S-acylated with both palmitate and stearate. HAP1 cells were labeled with palmitate (negative control, no alkyne group), alkyne-palmitate (Alk-C16), or alkyne-stearate (Alk-C18) for 3 hours followed by lysis and click chemistry. S-acyl-protein levels (isolated by click; left panels) and total protein levels in parent lysates (right panels) were assessed by immunoblot with antibodies against PK, PGK, GAPDH, and Actin. Quantified data are shown on the right (N=3-5; all comparisons not significant unpaired t-test).

### GAPDH is S-acylated by ZDHHC5 and ZDHHC17

S-acylation is catalyzed by the ZDHHC family of protein S-acyltransferases (PATs) named for their cytosolic zinc finger domain and highly conserved Asp-His-His-Cys (DHHC) catalytic motif^18,26^. Thus, we next sought to identify the GAPDH PAT(s). We focused on GAPDH as it is the first enzyme of the payoff phase of glycolysis, can be S-acylated *in vitro*^21^, and is the most well studied in the context of fueling fast axonal transport^16,17^. Mammals have 23 to 24 ZDHHC enzymes (ZDHHC1-24/25, skipping 10)^18,26^ so identifying the PAT can be a labour-intensive process. However, as GAPDH is largely a soluble protein that predominantly localizes in a diffuse manner throughout the cytosol (Figure S1), co-expression with a PAT that increases its S-acylation should result in increased association with membranes. As such, we used an imaging overexpression screen to identify candidate PATs^27,28^. HAP1 cells were co-transfected to express N-terminally GFP (green fluorescent protein)-tagged GAPDH and each of the 24 N-terminally HA (hemagglutinin)-tagged mouse ZDHHC enzymes individually. GAPDH localization was assessed by immunocytochemistry. Co-expression of GFP-GAPDH with either HA-ZDHHC5 or HA-ZDHHC17 resulted in the greatest change in localization of GFP-GAPDH from diffuse to punctate (Figure S1), suggesting that these two ZDHHC enzymes may S-acylate GAPDH.

To validate our findings from the imaging screen, we generated *ZDHHC5* or *ZDHHC17* knockout HAP1 cells using a CRISPR-Cas9 dual guide RNA (gRNA) strategy^29^. We verified loss of ZDHHC5 and 17 in the respective knockout cells by immunoblot (Figure 3A) and assessed GAPDH S-acylation in both cell lines compared to a control non-targeting (NT) cell line (Figure 3B) by ABE. GAPDH S-acylation was significantly reduced in ZDHHC17-deficient cells, with a non-statistically significant reduction in the ZDHHC5-deficient cells. S-acylation of Ras, a well-known substrate of a different PAT, ZDHHC9^30^, was readily detected in all three cell lines and remained unchanged with loss of ZDHHC17 or ZDHHC5 (Figure 3B). We also generated ZDHHC13 and ZDHHC14 knockout cells as controls and verified knockout by immunoblot (Figure S2A). ZDHHC13 was chosen due to homology with ZDHHC17 and as it is the only other ankyrin-repeat domain-containing PAT^31^ and ZDHHC14 was chosen as a negative control as its overexpression did not lead to the formation of cellular GFP-GAPDH puncta in the imaging screen (Figure S1). GAPDH S-acylation was unchanged in both ZDHHC13 and ZDHHC14 knockout cell lines (Figure S2B). Collectively, these data suggest that GAPDH is S-acylated by ZDHHC5 and/or ZDHHC17 in HAP1 cells.

**Figure 3.**
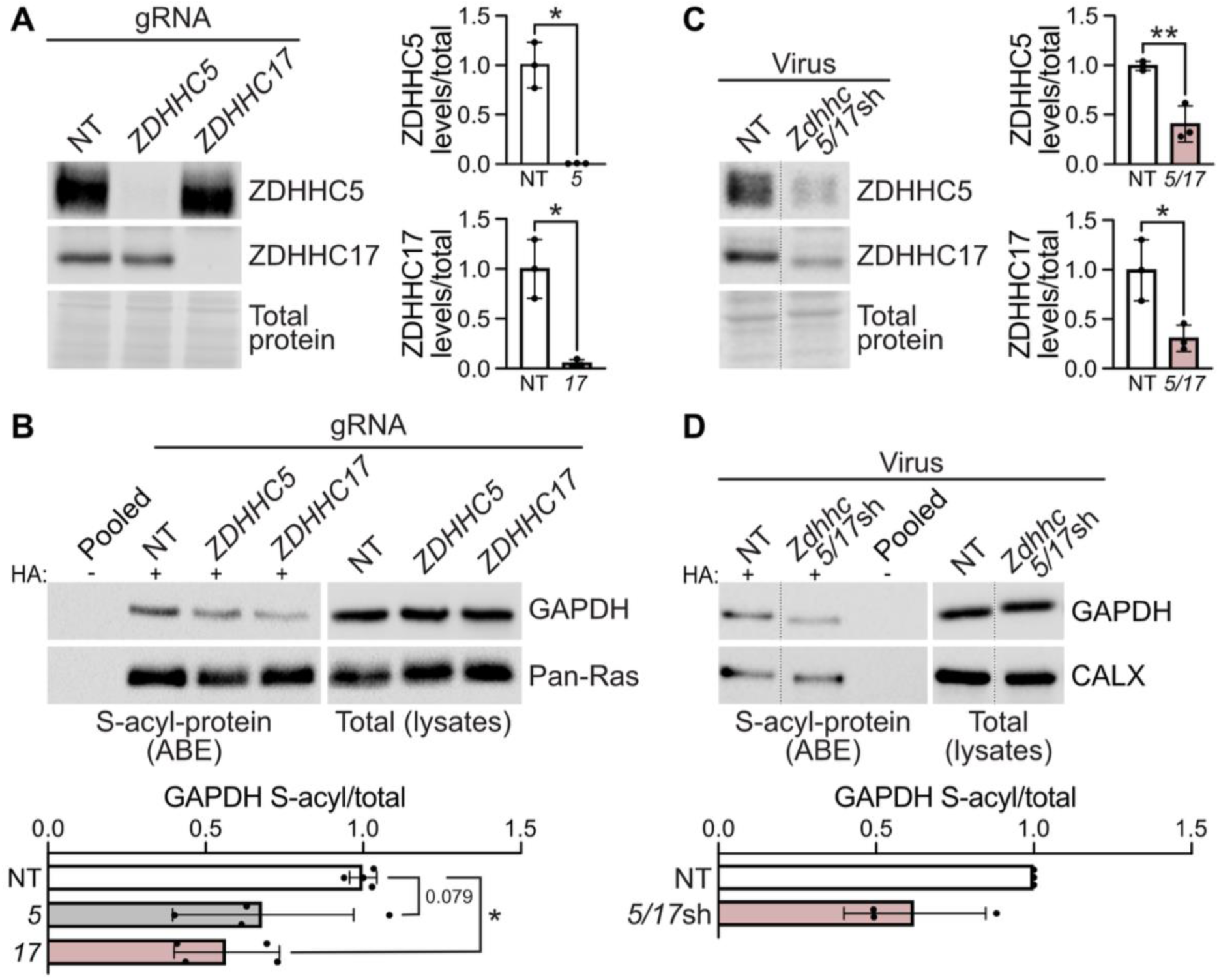
Loss of ZDHHC5 and/or ZDHHC17 reduces S-acylation of GAPDH. (**A**) ZDHHC5 and ZDHHC17 levels in lysates from NT (non-targeting gRNA), *ZDHHC5*, and *ZDHHC17* knockout HAP1 cells were determined by immunoblot using antibodies against ZDHHC5 (top panel) and ZDHHC17 (middle panel). Total protein levels using Ponceau S stain were determined (bottom panel). Quantified data are shown on the right with ZDHHC5 (top) or ZDHHC17 (bottom) levels relative to total protein, normalized to NT (ZDHHC5: Welch’s unpaired t test *p=0.018, N=3; ZDHHC17: Welch’s unpaired t test *p=0.030, N=3). (**B**) Cells were lysed and S-acyl-protein levels (isolated by ABE; left panels) and total protein levels in parent lysates (right panels) were assessed by immunoblot with antibodies against GAPDH and the positive control protein Ras (pan-Ras antibody). Quantified data are shown on the bottom with S-acyl/total levels, normalized to NT (one-way ANOVA with Dunnett’s multiple comparisons test, p=0.029, F(2,9)=5.39, *p<0.05, N=4). (**C**) ZDHHC5 and ZDHHC17 levels in lysates from hippocampal neurons transduced to express NT shRNA or shRNAs targeting *Zdhhc5* and *Zdhhc17* were determined by immunoblot using antibodies against ZDHHC5 (top panel) and ZDHHC17 (middle panel). Total protein levels were determined using Ponceau S (bottom panel). Quantified data are shown on the right with ZDHHC5 (top) or ZDHHC17 (bottom) levels relative to total protein, normalized to NT (ZDHHC5: unpaired t test **p=0.0058, N=3; ZDHHC17: unpaired t test *p=0.024, N=3). (**D**) Neurons were lysed and S-acyl-protein levels (isolated by ABE; left panels) and total protein levels in parent lysates (right panels) were assessed by immunoblot with antibodies against GAPDH and the positive control protein CALX. Quantified data are shown on the bottom with S-acyl/total levels, normalized to NT (N=3, not significant). Representative images in C and D are composites from the same immunoblot image, cuts are indicated by a dashed line. Pooled is a combined sample processed without the key ABE reagent HA (negative control).

To determine whether ZDHHC5 and ZDHHC17 are also PATs for GAPDH in neurons, we knocked down both enzymes/PATs (simultaneously) in primary rat hippocampal neurons using lentiviral delivery of short hairpin RNAs (shRNA; Figure 3C). Loss of ZDHHC5 and ZDHHC17 resulted in a trend to reduced GAPDH S-acylation (Figure 3D), suggesting that ZDHHC5 and ZDHHC17 are GAPDH PATs in neurons.

### GAPDH is S-acylated at cysteine 247

Next, we aimed to determine if blocking GAPDH S-acylation impairs binding to vesicles. To generate S-acyl-deficient GAPDH, we sought to identify GAPDH S-acylation site. The site of human GAPDH S-acylation is unknown, however, rabbit GAPDH is S-acylated at cysteine 244 *in vitro*^21^. It is thus likely that human GAPDH is S-acylated at the same cysteine, which is conserved in rabbit (*Oryctolagus cuniculus*), human (*Homo sapiens*), mouse (*Mus musculus*), and rat (*Rattus norvegicus*) but not in Zebrafish (*Danio rerio*), or fruit fly (*Drosophila melanogaster*; Figure 4A). To confirm the site of S-acylation we transfected HAP1 cells to express WT HA-GAPDH or a cysteine 247 (the homologous cysteine in human GAPDH) to serine (CS) variant and assessed S-acylation using ABE. Indeed, changing cysteine 247 to a serine significantly decreased GAPDH S-acylation (Figure 4B), suggesting that this is the predominant GAPDH S-acylation site.

**Figure 4.**
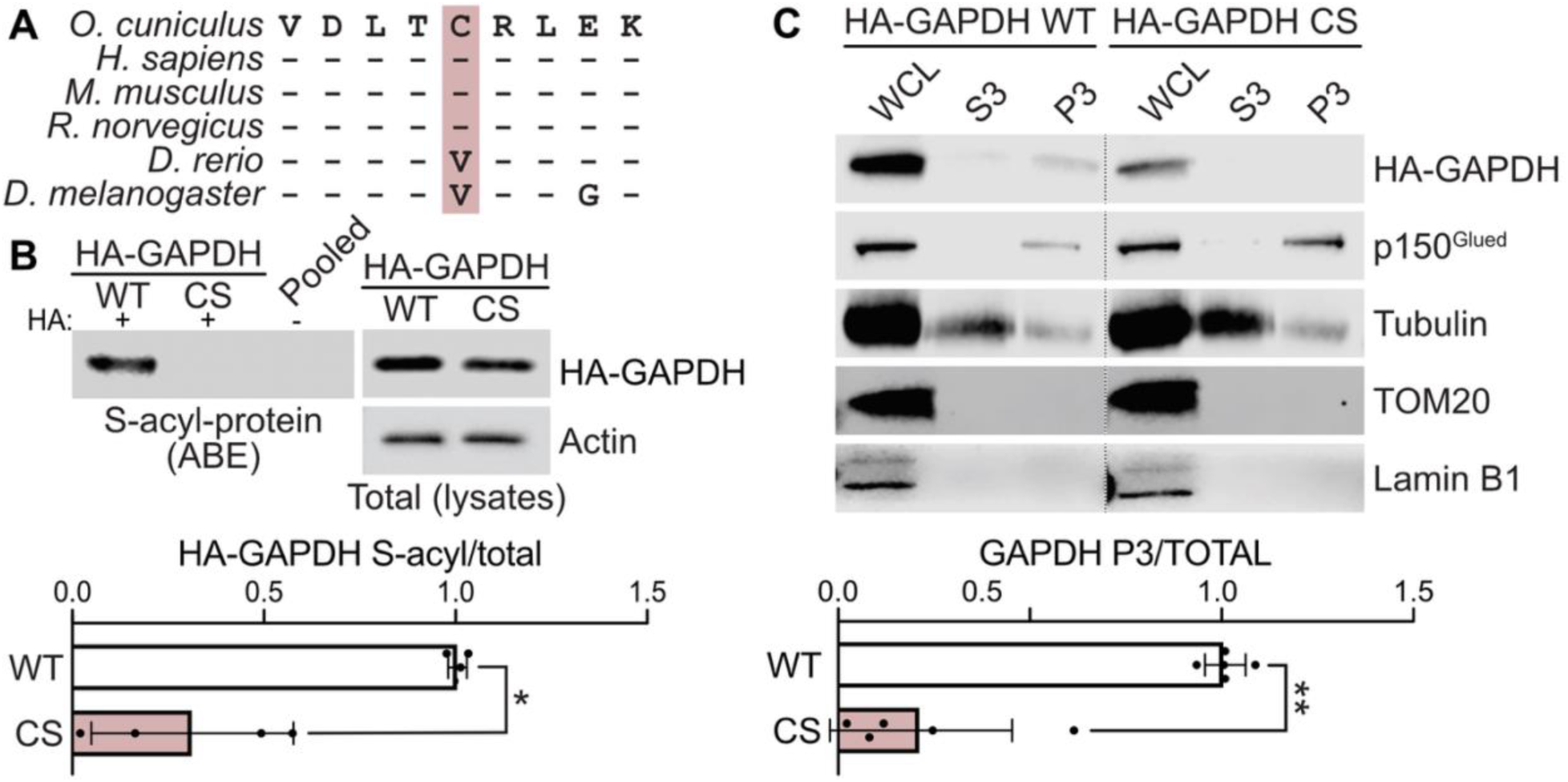
S-acylation of GAPDH at cysteine 247 is required for its association with the vesicle fraction in neurons. (**A**) Alignment of GAPDH amino acid sequence surrounding the putative S-acylation site from rabbit (*O. cuniculus*), human (*H. sapiens*), mouse (*M. musculus*), rat (*R. norvegicus*), zebrafish (*D. rerio*), and fruit fly (*D. melanogaster*). (**B**) HAP1 cells were transfected to express HA-tagged WT or a Cys (C) to Ser (S) GAPDH variant (C247S; CS). S-acyl-protein levels (isolated by ABE) and total protein levels in parent lysates were assessed by immunoblot with HA and Actin antibodies. Quantified data are shown below with S-acyl/total levels, normalized to WT (unpaired t-test with Welch’s correction, N=4, *p=0.013). Pooled is a combined sample processed without the key ABE reagent, HA (negative control). (**C**) Hippocampal neurons were transfected to express HA-tagged WT or CS GAPDH. Vesicle-associated proteins (P3 fraction) and cytosolic proteins (S3 fraction) were isolated by subcellular fractionation as well as total proteins from parent lysates (whole cell lysate [WCL]). GAPDH levels in all three fractions were then assessed with an HA antibody. The control proteins p150^Glued^ (vesicles), acetylated tubulin (microtubules), TOM20 (mitochondria), and Lamin B1 (nuclei) were also assessed. Quantified data are shown below with GAPDH in the P3 fraction relative to GAPDH in the WCL, normalized to WT (unpaired t-test with Welch’s correction, N=5, **p=0.0013). Representative images in C are composites from the same immunoblot image, cuts are indicated by a dashed line.

### Reduced association of S-acylation-deficient GAPDH with vesicles

To determine if S-acylation tethers GAPDH to vesicles in neurons, primary rat hippocampal neurons were transfected to express HA tagged WT or CS GAPDH and vesicles were isolated using subcellular fractionation. WT HA-GAPDH was detected in the vesicle fraction (P3), but S-acylation deficient HA-GAPDH CS was not (Figure 4C). Importantly, the vesicular fraction was positive for the vesicle-associated protein p150^Glued^, a subunit of the dynein-dynactin complex, but not for mitochondrial protein TOM20 or nuclear protein Lamin B1 (Figure 4C), confirming the fidelity of fractionation.

We next verified the fractionation results by assessing WT and CS HA-GAPDH localization to vesicles in neurons. Neurons were transfected to express WT or CS HA-GAPDH and brain-derived neurotrophic factor (BDNF)-mCherry to identify a subset of axonal transport vesicles, and GFP as a soluble cell fill. As GAPDH is primarily a soluble, cytosolic protein, it was localized in a diffuse manner throughout the neuron (Figure S3). This diffuse localization obstructs clear visualization of membrane-associated GAPDH (Figure S3). To address this issue, neuron cellular membranes were permeabilized with 0.01% saponin prior to fixation to remove all cytosolic proteins, allowing better visualization of the membrane localized GAPDH (Figure S3 & 5). The effectiveness of the saponin treatment was confirmed by the loss of GFP signal following permeabilization. Consistent with our biochemical results, the saponin permeabilization reduced the association of S-acylation deficient CS HA-GAPDH with vesicle-like structures while the S-acylation competent WT HA-GAPDH remained associated with these structures (Figure 5).

**Figure 5.**
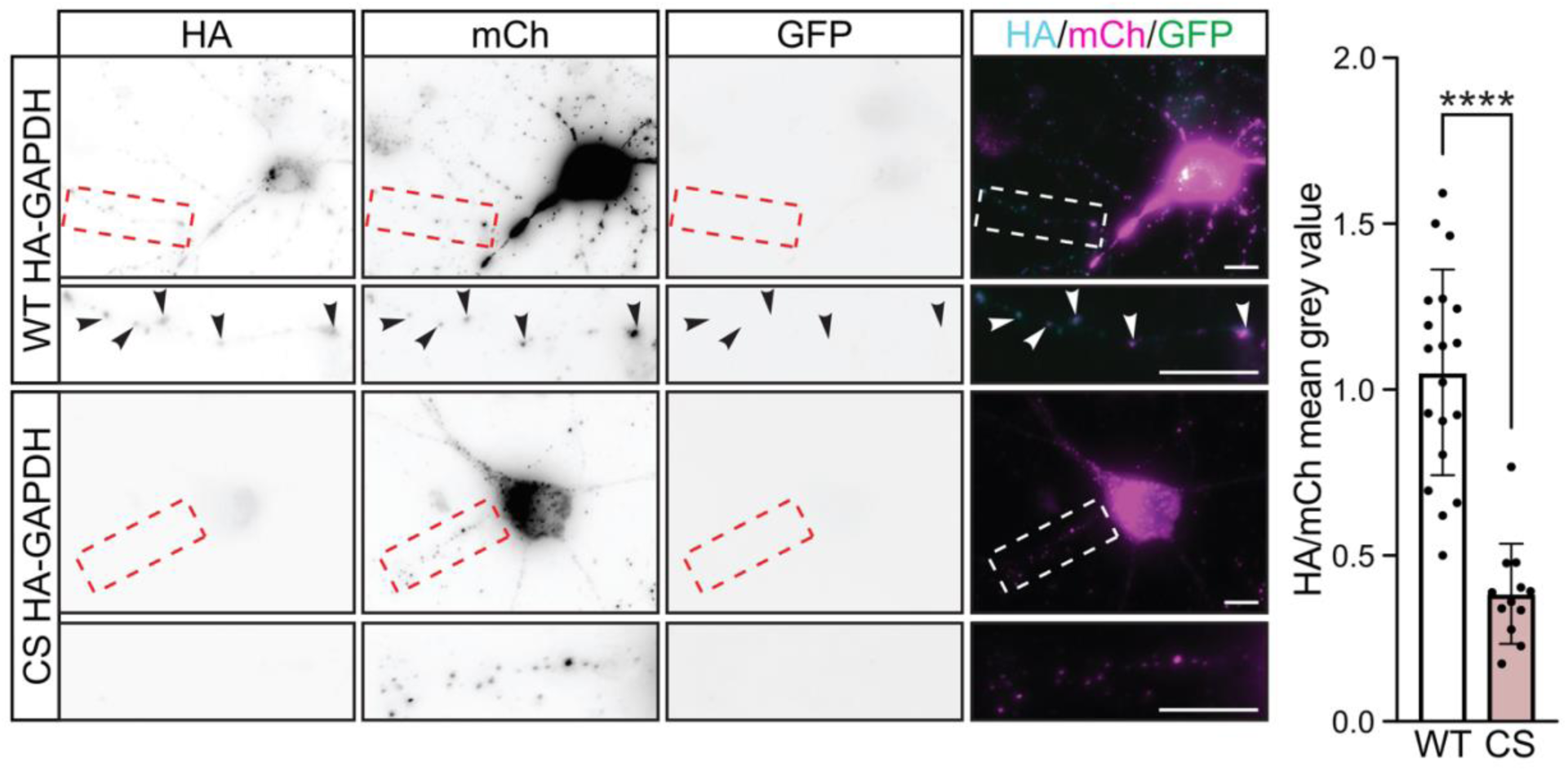
GAPDH S-acylation is required for its association with vesicles. Hippocampal neurons were co-transfected to express HA-tagged WT or CS GAPDH, BDNF-mCherry, and GFP. Prior to fixation, cells were permeabilized with 0.01% saponin for 5 minutes to remove cytosolic proteins. Cells were then fixed and immunostained with antibodies against HA (left panels; cyan in merged), RFP (to detect mCherry [mCh] second column panels; magenta in merged), or GFP (third column panels; green in merged). Lower panels show magnified view of red or white dashed box area in upper panels. Black arrowhead indicates puncta of GAPDH overlapped with BDNF. Scale bars are 10 μm. Quantified data are shown on the right with the mean grey value across the image for HA relative to that of mCh, normalized to WT (unpaired t-test, N=12-19 transfected neurons from two independent cultures, ****p<0.0001).

## Discussion

Our successful identification and low-throughput validation of glycolytic enzyme S-acylation provide a significant advancement towards elucidating the mechanisms that tether these enzymes to fast axonal transport vesicles. Here we show that eight out of ten glycolytic enzymes are S-acylated, and notably, that six are more S-acylated in the brain compared to the heart or kidney (Figure 1). This suggests that the S-acylation of glycolytic enzymes may facilitate a function specific to or of particular importance in the nervous system. One intriguing possibility is that this function is the tethering of the glycolytic enzymes to transport vesicles to fuel long-range transport, but this would require a more exhaustive cell type exploration *in vivo*. We also identified the human GAPDH S-acylation site, which allowed us to test whether S-acylation of GAPDH tethers it to vesicles. Both biochemical fractionation and immunocytochemistry data provide strong support for this hypothesis (Figures 4 and 5). Finally, we also found that GAPDH is S-acylated by ZDHHC5 and/or 17 (Figure 3).

The identification of ZDHHC5 and/or ZDHHC17 as GAPDH S-acylating enzymes is intriguing due to their potential roles in human neurological disorders. ZDHHC5 has been implicated in schizophrenia and ZDHHC17 in HD, amyotrophic lateral sclerosis, and in axon survival and degeneration^32–36^. ZDHHC17 is particularly compelling in the context of HD. ZDHHC17 enzymatic activity is impaired by mutant HTT in HD, which results in reduced S-acylation of ZDHHC17 substrates, including HTT itself^33,37–39^. Furthermore, fast axonal transport is defective in HD and restoring S-acylation by inhibiting the de-S-acylating enzyme acyl protein thioesterase 1 (APT1) restores trafficking in HD cell models^40^. Thus, it may be that impaired ZDHHC17 activity leads to reduced GAPDH S-acylation in HD thereby decreasing GAPDH tethering to fast axonal transport vesicles, contributing to fast axonal transport deficits. Notably, while both enzymes are localized to the Golgi, ZDHHC5 is also localized to the plasma membrane^41^. This may suggest that these enzymes are responsible for S-acylating different subcellularly localized pools of GAPDH, potentially in different contexts.

While we demonstrated that ZDHHC5 and/or ZDHHC17 S-acylate GAPDH, we observed only modest decreases in GAPDH S-acylation with loss of either ZDHHC5 or ZDHHC17 in HAP1 cells (30% and 45%, respectively, not significant for ZDHHC5) and with loss of both in neurons (40%; Figure 3). These modest effects may suggest the involvement of additional unidentified S-acylating enzyme(s) or that loss of both ZDHHC5 and ZDHHC17 is required to fully block S-acylation. Alternatively, insufficient double knockdown efficiency in neurons may account for the modest reduction observed in Figure 3D. Further exploration using higher knockdown efficiency or ZDHHC5/ZDHHC17 knockout neurons is required to confirm our findings.^41^

Association of S-acylation deficient HA-GAPDH with vesicles and vesicle-like structures is significantly reduced as measured by biochemical fractionation and immunocytochemical readouts (Figures 4 and 5). These data provide clear evidence that GAPDH S-acylation contributes to its tethering to vesicles in neurons. The incomplete block of association with vesicles and vesicle-like structures in our experiments may be due to overexpressed S-acylation deficient HA-GAPDH interacting with S-acylation competent endogenous GAPDH or with other glycolytic enzymes present on the vesicles. Indeed, GAPDH forms a tetramer^42^ so endogenous (S-acylation-competent) GAPDH may complex with S-acylation deficient GAPDH to facilitate binding to vesicles. Replicating our findings by knocking down endogenous GAPDH and replacing it with the S-acylation deficient variant will be important for determining whether this is the case. Alternatively, HTT contributes to scaffolding GAPDH on transport vesicles^17^, so exploring the interaction between GAPDH S-acylation and HTT in tethering GAPDH to vesicles is a critical area of future investigation. Also, HTT itself is an S-acylated protein, so exploring the importance of S-acylation of both proteins in their interaction with each other and with transport vesicles merits further investigation.

Interestingly, glycolytic enzymes have also been shown to form glycolytic complexes, termed metabolons or glycolytic bodies, in various cell types. These are associated with membranes and coupled to high-ATP demanding processes, including red blood cell membranes to fuel the Na^+^/K^+^ ATPase pump, neuronal synapses to support synaptic activity, and axonal growth cones to support growth^43–48^. Future work should explore the role of GAPDH S-acylation metabolon formation and determine whether other S-acylated glycolytic enzymes are similarly tethered to vesicles via this modification. Furthermore, it would be important to investigate whether all or only some component enzymes require S-acylation for inclusion in and attachment of the whole glycolytic enzyme complex to transport vesicles. This may also help explain why not all glycolytic enzymes are S-acylated in our experiments (Figure 1), as perhaps S-acylation of only a subset of metabolon components is sufficient to tether the entire complex.

While we have found that S-acylation tethers GAPDH to transport vesicles, it remains unknown how this process contributes to fast axonal transport. Assessing axonal transport dynamics by tracking fluorescently labelled cargo in the antero- and retrograde directions in live axons in GAPDH knockdown and replacement neurons, as well as *in vivo* experiments with WT or CS variants are thus key follow-up directions.

Our study is the first, to our knowledge, to provide evidence that S-acylation of GAPDH and other glycolytic enzymes may be contributing to their vesicle tethering in neurons and thus axonal transport. Efficient transport is vital for physiological function, neurodevelopment, and complex behaviors such as learning and memory. Autophagosomes, lysosomes, neurotrophins, signaling endosomes, and synaptic vesicles all rely on fast axonal transport. Our findings thus have direct implications for the neurodegenerative disease HD and likely broader implications for other disorders.

## Methods

All gRNA and shRNA sequences, as well as antibody information can be found in the Key Resources Table in the Supplementary Material.

### Animals

The use of animals was performed according to guidelines set by the Canadian Council on Animal Care and was approved by the Animal Care Committee of the University of Guelph (animal use protocols #4478 and #5320). Female timed pregnant adult Sprague-Dawley rats were purchased (Charles River Laboratories, Garfield Heights, OH, USA) and housed at the University of Guelph Central Animal Facility under a 12-hour light-dark cycle, with food and water available *ad libitum*. Rats were euthanatized using CO_2_ asphyxiation followed by decapitation. Hippocampal tissue was collected from day 18 embryos (E18) for primary cultures described below and the brain, heart, and kidney from the dam were harvested and flash frozen in liquid nitrogen and stored at -80°C until processing for ABE assay, as described below. As a result, all tissues were from female rats.

### Molecular Biology

Human *GAPDH* cDNA was purchased from the DNASU plasmid repository (clone HsCD00004782)^49^ and subcloned into the pHA-FEW or pGFP-FEW plasmids downstream of the human eukaryotic elongation factor 1a [EF1α] promotor at the XhoI/NotI sites to tag with HA or GFP, respectively, on the N-terminus^50,51^. A mutation was introduced in the *GAPDH* cDNA sequence to generate the C247S variant using the Q5 site-directed mutagenesis kit and primers designed with the NEBaseChanger Tool (https://nebasechanger.neb.com/; New England Biolabs (NEB), Ipswich, MA, USA). Pre-pro rat *Bdnf* (Brain Derived Neurotrophic Factor) cDNA was synthesized as a gene fragment (gBlock) with restriction enzyme sites by Integrated DNA Technology (IDT, Coralville, IA, USA) and cloned into the XhoI/NotI sites of pmCh-FEW^50^ to tag with mCherry on the N-terminus downstream of the EF1α promoter. An enhanced Blue Fluorescent Protein (EBFP2) mammalian expression construct was generated by PCR cloning eBFP cDNA from EBFP2-CathepsinB-6 (a gift from Michael Davidson (Addgene plasmid # 55236)^52^ into the BamHI and XbaI sites of pFEGW^53^ downstream of the EF1α promoter to replace GFP and generate pFEBW. Cassettes containing previously validated rat *Zdhhc5* or *Zdhhc17*-targeting shRNA sequences^34,54^ under the human U6 RNA polymerase III promoter were purchased from Genewiz as a custom gene synthesis plasmid (Genewiz, South Plainfield, NJ, USA) and inserted into the NheI and PacI sites of pFEBW. A previously published non-targeting shRNA sequence against luciferase that does not target any mammalian gene^55^ was cloned into pFEBW in the same way, but the cassette was purchased as a gene-block from Twist Biosciences (Twist Biosciences, San Francisco, CA, USA). N-terminally HA-tagged mouse *Zdhhc* cDNA plasmids with EF1ɑ promoter were previously described^53^. LentiCas9-Blast was a gift from Dr. Feng Zhang (Addgene plasmid #52962)^56^ and plasmids with dual CRISPR/Cas9 guide RNAs (gRNA; pCLIP-DUAL-SFFV-ZsGreen-sgRNA) targeting human *ZDHHC5*, *ZDHHC13*, *ZDHHC14*, *ZDHHC17*, or *eGFP* (non-targeting [NT], negative control)^29^ were purchased from the University of Ottawa GEM Facility. pRSV-Rev (Addgene plasmid # 12253), pMDLg/pRRE (Addgene plasmid # 12251), and pMD2.G (Addgene plasmid # 12259) were gifts from Dr. Didier Trono^57^. All other cloning reagents were from NEB, and constructs were verified by Sanger or nanopore sequencing.

### HAP1 cell culture

HAP1 cells were a kind gift from Dr. Leonardo Susta (University of Guelph) and were originally purchased from Horizon Discovery (Waterbeach, UK, #C631) and were cultured in Iscove’s Modified Dulbecco’s Medium (IMDM; Wisent Bioproducts, St. Bruno, QC, Canada, #319-105-CL), supplemented with 10% fetal bovine serum (FBS; Gibco, Waltham, MA, USA, #12483020), 1% penicillin/streptomycin (ThermoFisher Scientific, Waltham, MA, USA; #15140122), and 1% L-glutamine (Wisent Bioproducts, #609-065-EL) at 37°C with 5% CO_2_. 80-90% confluent 6-cm plates seeded the day before were used for bioorthogonal labeling and click chemistry or transfection and ABE assay described below. HAP1 cells were plated on poly-L-lysine hydrobromide (PLL; Millipore Sigma, Burlington, MA, USA, #P2636) coated glass coverslips for transfection and immunocytochemistry described below.

### HAP1 cell transfection

HAP1 cells were transfected using a previously described polyethylenimine (PEI) transfection (Millipore Sigma, #919012)^58,59^. A PEI-DNA-IMDM mixture with a 1:5 ratio of PEI:DNA was made by mixing 5x volume (of the DNA μg amount) of 1mg/mL PEI with media and adding 1-3 μg of DNA. This mix was briefly vortexed and incubated for 10 minutes at room temperature prior to adding it dropwise to 80-90% confluent cells seeded the day before. Cells were recovered after six hours, and the following day were harvested for the ABE assay or fixed for immunocytochemistry as described below.

### Gene knockout in HAP1 cells

*ZDHHC5, ZDHHC13, ZDHHC14, or ZDHHC17* knockout HAP1 cells were generated as previously described^60^. 62,000 cells seeded in one well of a 6-well plate were transfected with 1.2 μg of Lenti-Cas9-BLAST and 0.8 μg of pCLIP-DUAL-sgRNA using PEI as described above. Cells were selected 48 hours later with 2 μg/mL puromycin (BioShop Canada Inc, Burlington, ON, Canada, #PUR333.25) until the cells in an untransfected well had all died. Following selection, cells were recovered in puromycin-free media for 7 days and were then maintained as a polyclonal population with a passage number-matched NT control line (*ZDHHC5* and *ZDHHC17* knockouts with a paired NT and *ZDHHC13* and *ZDHHC14* knockouts with a separate paired NT). 80-90% confluent cells seeded the day before were harvested for the ABE assay, as described below.

### Generation of lentivirus

HEK293T (human embryonic kidney) cells were cultured in complete standard Dulbecco’s Modified Eagle Medium (DMEM; Wisent Bioproducts; #319-015-CL) supplemented with 10% FBS, 1% penicillin/streptomycin, and 1% L-glutamine. Cells were seeded on 10-cm plates to achieve >90% confluency the following day in DMEM with FBS and L-glutamine as above but lacking antibiotics and transfected using an optimized CaPO_4_ protocol^54,61,62^ with a mix of 7.5 μg of DNA being packaged, 4.9 μg pMDLg/pRRE, 2.6 μg pMD2.G, and 1.9 μg pRSV-Rev. The DNA was mixed and incubated for 10 minutes, diluted in 500 μL 244 mM CaCl_2_, and combined dropwise with 500 μL pre-warmed, 37°C, 2x HEPES (4-(2-hydroxyethyl)-1-piperazineethanesulfonic acid) buffered saline (270 mM NaCl, 1.5 mM Na_2_HPO_4͘_•7H_2_O, 40 mM HEPES, 10 mM KCl, 10 mM D-glucose, pH 7.0) while mixing. This mixture was incubated for one minute at 37°C before being added dropwise to the cells. The cells were recovered in complete DMEM after 6-8 hours. The next day, cells were treated with 10 mM sterile sodium butyrate (VWR, Radnor, PA, USA; #TCS0519) for 4-6 hours before being placed in Neurobasal medium (Gibco, #21103-049) with 1% penicillin/streptomycin, 1% GlutaMAX (Gibco, #35050061), and 0.1% heat inactivated FBS. 24 hours later, the media was collected and spun for 10 minutes at 500 x g to pellet cell debris. The supernatant was then filtered through a 0.45 μm filter, aliquoted, snap frozen in liquid nitrogen, and stored at -80°C until use.

### Hippocampal neuron culture

Hippocampal neurons were cultured from E18 rat hippocampi in Neurobasal medium with 1% GlutaMAX, 2% B-27 supplement (Gibco, #17504-044), and 1% penicillin/streptomycin at 37°C in 5% CO₂ as previously described^54^. On day *in vitro* (DIV) 5, neurons were treated with the mitotic inhibitor 0.625 μM 5-Fluoro-2’-deoxyuridine/uridine (Millipore Sigma, #2F0503-100MG and #6U3003-5G, respectively). Neurons were seeded at a density of 280,000 cells/well in 6-well plates and transfected or transduced as described below for biochemical assays. Neurons were also seeded at a density of 180,000 cells/well in 6-well plates on PLL coated glass coverslips for transfection and immunocytochemistry described below.

### Hippocampal neuron transfection and transduction

Dissociated hippocampal neurons were electroporated prior to plating using a Nucleofector^TM^ 2b device (Lonza Biosciences, Basel, Switzerland) using the Nucleofector® Rat Neuron Kit (Lonza Biosciences, #VPG-1003), according to the manufacturer’s instructions using program 003. Neurons were then plated and cultured as described above until harvesting for biochemical fractionation.

Hippocampal neurons on coverslips in 6 well plates were transfected with Lipofectamine 2000 (L2K; ThermoFisher Scientific, #11668027) on DIV 18. 250 ng of each DNA was combined and incubated at room temperature for five minutes prior to dilution in 50 μL of plain Neurobasal media. The DNA-Neurobasal mix was combined dropwise with an equal volume of Neurobasal with 3 μL of L2K. DNA-L2K complexes were incubated for 15 minutes at 37°C before being added dropwise to neurons. After 90 minutes, neurons were recovered into a 1:1 solution of conditioned media (removed prior to transfection) and fresh complete Neurobasal. Neurons were fixed as described below the following day.

Neurons were transduced on DIV 6 using lentivirus. Frozen viral aliquots were quick thawed and the appropriate volume of virus was added directly into the neuron media. After 16 hours, neurons were recovered into a 1:1 solution of conditioned media (removed prior to transduction) and fresh complete Neurobasal.

### Bioorthogonal labelling and click chemistry

Bioorthogonal labelling and click chemistry were performed as previously described^63^. HAP1 cells were metabolically labelled with 100 μM alkyne fatty acid analogues (Vector Laboratories, Newark, CA, USA, alkyne-palmitate [15-hexadecynoic acid] #1165 or alkyne stearate [17-octadecynoic acid] #1166) or 100 μM non-alkyne palmitic acid (Millipore Sigma, #P5585) for 3 hours. Prior to labeling, fatty acids were saponified and bound to fatty acid-free bovine serum albumin (BSA; Millipore Sigma, #1003054814). Cells were lysed in 500 μL of modified radioimmunoprecipitation (RIPA) buffer (1M HEPES pH 7.4, 150 mM NaCl, 0.5% sodium deoxycholate, 1% Igepal CA-630, 0.1% SDS (sodium dodecyl sulfate), 2 mM MgCl_2_, and Roche *cOmplete™, EDTA-free Protease Inhibitor Cocktail* [PIC; Millipore Sigma, #11873580001]) with rotation at 4°C for 30 minutes, centrifuged at 13,000 revolutions per minute for 10 minutes, and filtered through 0.22 μm Spin-X^®^ centrifuge filters (Millipore Sigma, #CLS8161). Protein concentrations were determined using the *DC* (Detergent Compatible) Protein Assay (Bio-Rad Laboratories, Hercules, CA, USA, #5000111) according to the manufacturer’s instructions.

150-500 μg of lysate was added to 5 mM of BTTAA (2-(4-((Bis((1-(*tert*-butyl)-1*H*-1,2,3-triazol-4-yl)methyl)amino)methyl)-1*H*-1,2,3-triazol-1-yl)acetic acid; Vector Laboratories, #CCT-1236-100), 12.45 mM Sodium-L-ascorbic acid (Milipore Sigma, #A7631), 2.4 mM CuSO_4_·5H_2_O (Milipore Sigma, #209198-100G), and 100 μM Biotin Azide Plus (Vector Laboratories, #CCT-1488-25). The click reaction was performed for one hour in the dark at room temperature, then stopped with 10 mM EDTA (ethylenediaminetetraacetic acid). The protein was precipitated by adding 80% ice cold acetone overnight at -20°C and protein pellets were dissolved in ABE lysis buffer (50 mM HEPES pH 7.0, 1 mM EDTA, 2% SDS, and PIC) and diluted in dilution buffer (50 mM HEPES pH 7.0, 1% Triton X-100, 1 mM EDTA, 1 mM EGTA [ethylene glycol-bis(β-aminoethyl ether)-*N*,*N*,*N*′,*N*′-tetraacetic acid], 2.1 μM Leupeptin [Bioshop, #LEU001.25], and 1 mM Benzamidine [Bioshop, BEN666.25]). Biotinylated proteins were affinity-purified using Pierce high-capacity NeutrAvidin agarose beads (ThermoFisher Scientific, #PI29204) at 4°C for 3 hours. Subsequently, beads were washed 3 times using dilution buffer with 0.5 M NaCl and once with dilution buffer without NaCl. Purified S-acyl proteins were eluted using a hydroxylamine-based elution buffer (50 mM HEPES pH 7.4, 150 mM NaCl, 0.1% SDS, 1 M hydroxylamine hydrocholoride [HA; Millipore Sigma, #159417-500G], and PIC) by shaking at 21°C for one hour. Finally, the eluted protein was denatured in sample loading buffer (2% SDS, 50 mM Tris pH 6.8, 10% glycerol, 0.05% bromophenol blue, 1% β-mercaptoethanol [βME]) by heating for 5 minutes at 95°C. In addition, 25 μg of parent lysates was diluted in 100 μL of RIPA buffer with sample loading buffer and denatured as above. Eluted and lysate samples were subjected to SDS-PAGE (polyacrylamide gel electrophoresis) and immunoblot as described below.

### Tissue lysis

Frozen rat tissue samples were homogenized on ice in 1 mL/100 mg tissue homogenization buffer (10 mM phosphate buffer pH 7.4, 0.32 M sucrose, 1 mM EDTA, 8 M Urea, and PIC) with 100 mM N-ethylmaleimide (NEM; Millipore Sigma, #E3876). Proteins were then solubilized with a final concentration of 2.5% SDS (Fisher Bioreagents, Pittsburgh, PA, USA, #BP166-500) and DNA was sheared by sonicating 5 times for 6 seconds at 25% power. Cleared lysates were used for the ABE assay as described below.

### ABE assay

The ABE assay was performed as previously described^54^. HAP1 cells or neurons were lysed in ABE lysis buffer with 100 mM NEM. DNA was sheared by sonicating 4 times for 6 seconds at 20-25% power. Cleared tissue or cell lysates were incubated at 50°C with shaking for 60 minutes. Protein was precipitated as described above and protein pellets were dissolved in 4% SDS buffer (50 mM Tris pH 7.5, 5 mM EDTA, 4% SDS, and PIC). Protein concentrations were determined as described above. 250-1500 μg of protein was incubated rotating in the dark for one hour at room temperature with 1 mM Biotin-HPDP (APExBIO Technology LLC, Houston, TX, USA, #A8008) plus either 0.7 M hydroxylamine or 50 mM Tris pH 7.4 (HA-, a negative control with sample from each group pooled). Protein was precipitated again, as above, and protein pellets were dissolved in ABE lysis buffer (without NEM), diluted in dilution buffer, and purified as described above. Purified S-acyl proteins were eluted using 1% βME in 40 μL of elution buffer (0.2% SDS and 250 mM NaCl in dilution buffer) at 37°C for 10 minutes. Finally, the eluted protein was denatured in sample loading buffer as described above. In addition, 25-95 μg of parent lysates was diluted in 100 μL of RIPA buffer with sample loading buffer and denatured as above. Eluted and lysate samples were subjected to SDS-PAGE and immunoblot as described below.

### Biochemical subcellular fractionation

Vesicle purification was based on a previously established protocol^64^. Electroporated neurons were harvested on DIV 19 by scraping into ice-cold artificial cerebral spinal fluid (aCSF; 25 mM HEPES pH 7.4, 120 mM NaCl, 5 mM KCl, 2 mM CaCl_2_, 20 mM glucose, and 1 mM MgCl_2_) and collected by centrifugation at 2000 x g. The neuron pellet was immediately frozen on dry ice and stored at -80°C until processing. Neurons were ground up using a pestle, resuspended in 500 μL of homogenization buffer (320 mM sucrose, 1 mM EDTA, 4 mM HEPES pH 7.3 and PIC), and fully homogenized using a Dounce homogenizer. 40 μL was removed and lysed in RIPA buffer as the whole cell lysate (WCL) and the remainder was centrifuged for 10 min at 47,000 x g at 4°C (Beckmann MLA-130 rotor). The supernatant (S1) was collected, and the pellet was washed with 100 μL homogenization buffer and re-centrifuged and the supernatant pooled with the first collection. The pellet was resuspended with 75 μL of RIPA buffer forming the P1 fraction containing large cell fragments and nuclei. 40 μL of S1 was removed and lysed in RIPA buffer and the remaining was centrifuged for 40 min at 120,000 x g at 4°C. The pellet was re-suspended with 55.5 μL of RIPA buffer forming the P2 fraction containing large membrane fragments. 40 μL of the S2 supernatant fraction containing vesicles and other cell fragments was removed and lysed in RIPA buffer and the remaining was transferred onto a 100 μL sucrose cushion (700 mM sucrose and 10 mM HEPES pH 7.3) and centrifuged for 2 hours at 260,000 x g at 4°C. The pellet containing purified vesicles (P3) was resuspended in 55.5 μL of RIPA buffer, and the supernatant formed the S3 fraction. The samples were then denatured in sample loading buffer and 50 μL of each fraction was subjected to SDS-PAGE and immunoblot as described below.

### Immunoblot analysis

Samples prepared as described above were subjected to SDS-PAGE. Gels were transferred onto 0.45 μm nitrocellulose membranes (Bio-Rad Laboratories, #1620115) and blocked with 5% skim milk/Tris-buffered saline (TBS) prior to immunoblotting. Primary antibody dilutions in 1% BSA (Millipore Sigma, #A-3912)/TBS were applied to membranes and incubated at 4°C overnight. Membranes were washed 3 times for 5 minutes in TBS with 0.5% tween-20 (TBST) prior to incubation with the corresponding fluorescent- or HRP (horse radish peroxidase)-conjugated secondary antibodies in 5% milk/TBS at 1:5000 for 1 hour at room temperature. Membranes were then washed 4 times 5 minutes in TBST and HRP-conjugated secondary antibody signal was detected using Clarity Western ECL Substrate (Bio-Rad Laboratories, #1705061). Chemiluminescence and fluorescence signals were imaged with the Odyssey Fc Imaging System (Li-COR Bioscience, Lincoln, NE, USA) or the ChemiDoc XRS+ with Image Lab Software (Bio-Rad Laboratories) and quantified using Image Studio Software Version 6 (Li-COR Biotech, Lincoln, NE, USA). All antibodies and dilutions are listed in the Key Resource Table in Supplementary Information.

### Immunocytochemistry

Transfected hippocampal neurons were washed with aCSF prior to incubation in aCSF alone or aCSF with 0.01% saponin for 5 minutes. Neurons were immediately fixed in Parafix (4% paraformaldehyde [Electron Microscopy Sciences, Morgantown, PA, USA, #15714S] and 4% sucrose in phosphate buffered saline [PBS]) at 37°C for 30 min. Coverslips were then washed 3 times in PBS followed by blocking and permeabilization with 0.25% Triton X-100 and 0.2% gelatin in PBS for one hour. Coverslips were subsequently washed 3 times for five minutes with PBS prior to incubation with primary antibodies overnight at 4°C. The following day, coverslips were washed as above, incubated with secondary antibodies for one hour at room temperature, and washed again prior to mounting onto microscope slides using ProLong™ Gold Antifade mountant (ThermoFisher Scientific, #P36934). All antibodies and dilutions are listed in the Key Resource Table in Supplementary Information.

### Image acquisition and analysis

Immunostained HAP1 cells and neurons were imaged using the Nikon ECLIPSE T*i*2 fluorescent microscope with a 60x oil immersion objective (1.4 numerical aperture [NA], PL-APO).

Neurons were imaged using 0.4 μm spaced Z-stacks. Parameters were kept constant between coverslips and experiments. All neuron images are maximum intensity projections images modified for brightness and contrast using ImageJ Fiji^65^. Analysis of HA-GAPDH vesicle association was determined by measuring the mean grey value across the image for the HA channel using Fiji and was calculated relative to the mean grey value across the image for the BDNF-mCherry channel to account for neuron-to-neuron variation in transfection efficiency.

### Statistical analysis

All data was analyzed using Microsoft Excel (Microsoft, Redmond, WA, USA), and GraphPad Prism 11 software (GraphPad Software, San Diego, CA, USA). All graphs show mean with standard deviation of the mean. A Shapiro-Wilk normality test was performed to determine if the data were normally distributed and an F or Barlett’s test was used to determine equal variance. The statistical test used is indicated in all figure legends. Individual rats, HAP1 cells of different passage numbers, and neuron cultures established from separate litters were considered biological replicates.

