## Supplementary material for "GAPDH is tethered to axonal transport vesicles by S-acylation"

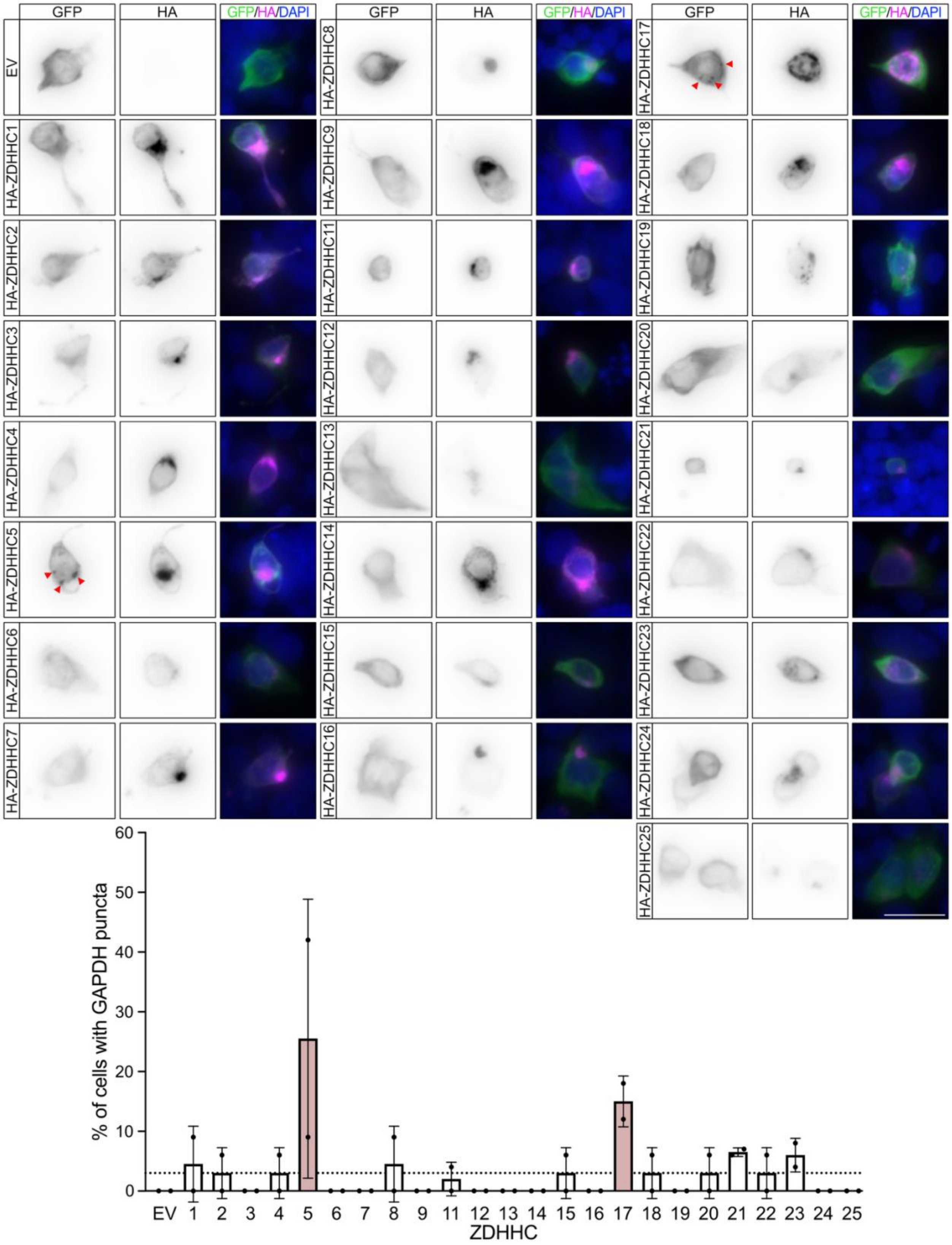


**Figure S1: Protein S-acyltransferases ZDHHC5 and ZDHHC17 are putative GAPDH S-acylating enzymes.** HAP1 cells were transfected to express GFP-GAPDH and each individual HA-tagged ZDHHC enzyme. Cells were fixed and immunostained with antibodies against GFP (left panels, green in merged) and HA (middle panels, magenta in merged) along with a DAPI DNA stain (blue in merged). The percentage of double transfected cells with GFP-GAPDH puncta, indicated by red arrowheads in left panels, for each ZDHHC enzyme is shown on the bottom (N=2). Dashed line in graph indicates the average value across all conditions. An empty vector (EV) was included as a negative control. Scale bar is 10 µm.

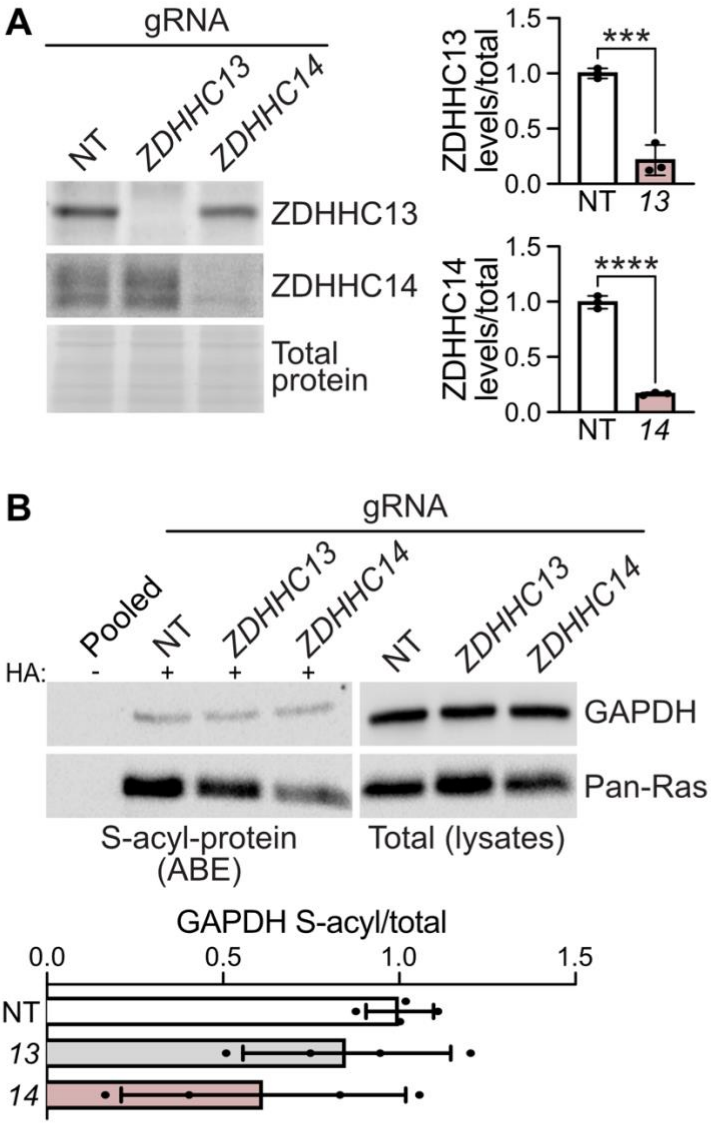


**Figure S2: Loss of ZDHHC13 and ZDHHC14 does not impair S-acylation of GAPDH.** (**A**) ZDHHC13 and ZDHHC14 levels in lysates from NT, *ZDHHC13*, and *ZDHHC14* knockout HAP1 cells were determined by immunoblot using antibodies against ZDHHC13 (top panel) and ZDHHC14 (middle panel). Total protein levels using Ponceau S stain were determined (bottom panel). Quantified data are shown on the right with ZDHHC13 (top) or ZDHHC14 (bottom) levels relative to total protein, normalized to NT (ZDHHC13: unpaired t test $p=0.0007$, N=3; ZDHHC14: unpaired t test $p<0.0001$, N=3). (**B**) Cells were lysed and S-acyl-protein levels (isolated by ABE; left panels) and total protein levels in parent lysates (right panels) were assessed by immunoblot with antibodies against GAPDH and the positive control protein pan-Ras. Quantified data are shown on the bottom with S-acyl/total levels, normalized to NT. Pooled is a combined sample processed without the key ABE reagent, hydroxylamine (HA; negative control).

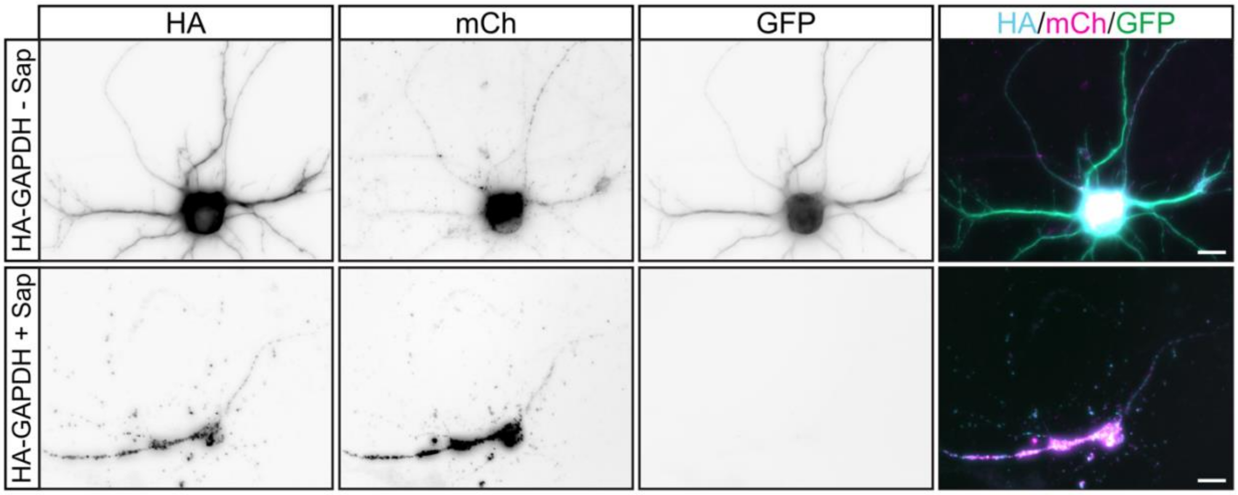


**Figure S3: HA-GAPDH is localized to the cytosol and vesicles in neurons.** Hippocampal neurons were co-transfected to express HA-tagged WT GAPDH, BDNF-mCherry, and GFP. Prior to fixation, cells were permeabilized with 0.01% saponin in aCSF (+ Sap) or treated with aCSF alone (- Sap) for 5 minutes to remove cytosolic proteins. Cells were then fixed and immunostained with antibodies against HA (left panels; cyan in merged), RFP (to detect mCherry [mCh] second column panels; magenta in merged), or GFP (third column panels; green in merged). Scale bar is 10 µm.

**Table 1.** Key Resources table.

| | | |
|---|---|---|
| **gRNA sequences** | | |
| **Gene** | **Sequence** | |
| *ZDHHC5* | gRNAa ACTGCAGTGGGAACATCGAG and gRNAb GGAACTCTCAGGGGTCCGCA | |
| *ZDHHC13* | gRNAa CCATGTATGCAACTGCTGTG and gRNAb GAGGTGGCTGCAGAAATGCG | |
| *ZDHHC14* | gRNAa AGAAGAAAATCGCGGCCCGG and gRNAb CTTTGGTTCTGGGAGGCGGG | |
| *ZDHHC17* | gRNAa AATTCTGGATGTATGTGACG and gRNAb GCTATTGTGGATCAACTTGG | |
| Non-targeting eGFP | gRNAa CGAGGAGCTGTTCACCGGGG and gRNAb CAACTACAAGACCCGCGCCG | |
| **shRNA sequences** | | |
| **Gene** | **Sequence** | |
| Non-targeting (luciferase) | CGTACGCGGAATACTTCGA | |
| *Zdhhc5* | CCTCAGATGATTCCAAGAGAT | |
| *Zdhhc17* | ATGAATGCCAGGAGATACAAGCACTTTAA | |
| **Antibodies** (IB=immunoblot; ICC=immunocytochemistry) | | |
| **Target, type, & dilution** | **Catalogue number** | **Manufacturer** |
| Anti-HA rabbit polyclonal; IB 1:5000 | #S3724 | Cell Signaling Technologies, Danvers, MA, USA |
| Anti-HA mouse monoclonal; ICC 1:500 | HA.11, #901514 | BioLegend, San Diego, CA, USA |
| Anti-GFP chicken polyclonal; ICC 1:500 | #AB16901 | Millipore Sigma |
| Anti-RFP (red fluorescent protein) rat | #5f8 | Proteintech, Rosemond, IL, USA |

| | | |
|---|---|---|
| monoclonal; ICC 1:1000 | | |
| Anti-TOM20 rabbit polyclonal; IB 1:2000 | #11802-1-AP | Proteintech |
| Anti-HK rabbit polyclonal; IB 1:1000 | #NBP1-90177 | Novus Biologicals, Centennial, CO, USA |
| Anti-GPI rabbit polyclonal; IB 1:1000 | #NBP1-90177 | Novus Biologicals |
| Anti-PFK rabbit polyclonal; IB 1:1000 | #13389-1 | Proteintech |
| Anti-ALDO rabbit polyclonal; IB 1:1000 | #A304-494A | Bethyl Laboratories, Montgomery, TX, USA |
| Anti-TPI rabbit polyclonal; IB 1:1000 | #A303-755A-M | Bethyl Laboratories |
| Anti-GAPDH rabbit polyclonal; IB: 1:50000 | #60004-1-Ig | Proteintech |
| Anti-PKM1/2 rabbit monoclonal; IB 1:1000 | #3190S | Cell Signaling Technologies |
| Anti-PGAM rabbit monoclonal; IB 1:1000 | #12098S | Cell Signaling Technologies |
| Anti-ENOL rabbit monoclonal; IB 1:1000 | #3810T | Cell Signaling Technologies |
| Anti-Calnexin rabbit monoclonal; IB 1:5000 | #10427-2-AP | Cell Signaling Technologies |
| Anti-CREB mouse monoclonal; IB 1:1000 | #9104S | Cell Signaling Technologies |
| Anti-Actin mouse monoclonal; IB 1:1000 | #3700T | Cell Signaling Technologies |
| Anti-p150Glued mouse monoclonal; 1:250 | #610473 | BD Biosciences, Franklin Lakes, NJ, USA |
| Anti-Acetylated Tubulin mouse monoclonal; IB 1:5000 | #T7451 | Millipore Sigma |
| Anti-Lamin B1 rabbit polyclonal; IB 1:5000 | #12987-1-AP | Protientech |
| Anti-ZDHHC13 rabbit polyclonal; IB 1:500 | #24759-1-AP | Proteintech Group, Inc, Rosemont, IL, USA |
| Anti-ZDHHC17 rabbit polyclonal; IB 1:1000 | #H7414 | Millipore Sigma |
| Anti-ZDHHC5 rabbit polyclonal; IB 1:1000 | #HPA014670 | Millipore Sigma |
| Anti-ZDHHC14 rabbit polyclonal; IB 1:200 | - | Sanders SS, et al, 2020, eLife |
| Anti-pan Ras rabbit polyclonal; IB 1:1000 | #MABS195 | Millipore Sigma |
| Alexa Fluor 568-conjugated goat anti-rat; ICC 1:500 | #A-11077 | ThermoFisher Scientific |
| Alexa Fluor 488-conjugated goat anti-chicken; ICC 1:500 | #A-11039 | ThermoFisher Scientific |
| Alexa Fluor 647-conjugated goat anti-mouse; ICC 1:500 | #A-21242 | ThermoFisher Scientific |
| HRP-conjugated donkey anti-rabbit; IB 1:5000 | #AP182P | MilliporeSigma |
| HRP-conjugated horse anti-mouse; IB 1:2000 | #7076 | Cell Signaling Technologies |
| Alexa Fluor 680-conjugated goat anti-mouse; IB 1:5000 | #A21057 | ThermoFisher Scientific |
| Alexa Fluor 680-conjugated rabbit anti-goat; IB 1:5000 | #A21088 | ThermoFisher Scientific |
| Alexa Fluor 680-conjugated donkey anti-rabbit; IB 1:5000 | #A10043 | ThermoFisher Scientific |
| IRDye 800-conjugated donkey anti-rabbit; IB 1:5000 | #926-32213 | Li-COR Biosciences |
